# Real-time analysis of pore formation by bi-component staphylococcal leukotoxins using the two-electrode voltage-clamp technique

**DOI:** 10.64898/2026.08.03.742423

**Authors:** Laura Lemel, Simon Harris, Guillaume Audic, Laure Bellard, Jean-Daniel Savoie, Claire M. Grison, Sébastien Granier, Justine Magnat, Normand Voyer, Thierry Vernet, Isabel D. Alves, Anne-Marie Di Guilmi, Christophe J. Moreau

**Affiliations:** Univ. Grenoble Alpes, CNRS, CEA, IBS, 71, av. Martyrs, CS10090, F-38044 Grenoble, France; Département de chimie, PROTEO and Centre d’études nordiques, Université Laval, Québec, QC, Canada G1V 0A6; IGF, University of Montpellier, CNRS, INSERM, Montpellier, France; Institut de Chimie et de Biologie des Membranes et des Nano-objets (CBMN), Université de Bordeaux, CNRS, UMR 5248, 33600 Pessac, France

## Abstract

Pore forming toxins (PFTs) are cytotoxins secreted in water-soluble form by pathogenic bacteria. They have the ability to form pores in the membrane of host cells, ultimately leading to cell death by lytic activity. *Staphylococcus aureus* produces a variety of bi-component PFTs, the leukocidins, which target and lyse particular leukocytes, erythrocytes and endothelial cells through specific interactions with membrane receptors. Most of these receptors belong to the family of complement or chemokine receptors that are G protein-coupled receptors (GPCRs). Gamma-hemolysins (Hlgs) are the major leukocidins secreted by *S. aureus*, and form receptor-dependent hetero-octameric pores through mechanisms that are not fully elucidated. Studying these molecular mechanisms is technically challenging due to the requirement of specific receptors in a lipid bilayer environment. In the present article, we developed a simple and highly sensitive method allowing cell surface expression of a large diversity of target receptors and recording in real-time, currents generated by neo-formed pores. This method is based on the heterologous expression of receptors in *Xenopus* oocytes and on the two-electrode voltage-clamp technique with electrophysiological robots. Using this approach, we characterized the concentration dependent-kinetics of pore formation, determined the receptor density as a limiting factor, showed specific response to non-cognate pairing of PFTs, observed cell surface binding of F subunits preceding pore formation and propose a hybrid model of subunit oligomerization. This method could be easily implemented for the *in vitro* characterization of various PFTs on a wide diversity of membrane receptors, to decipher early mechanisms of pore formation or to screen therapeutic agents blocking the cytotoxicity of receptor-dependent PFTs.

**Author Summary:** *Staphylococcus aureus* is a bacterial species naturally present in our external flora and environment, but it is also one of the main pathogens responsible for nosocomial infections in hospital, with strains having highly problematic multi-resistance to antibiotics. *S. aureus* is able to secrete various virulence factors, some of which can specifically target and lyse our immune cells, making us more vulnerable to this pathogen. Thus, leukotoxins bind to receptors on the cell surface, drastically change their conformation and form cytotoxic pores in the membrane. Studying the molecular mechanisms underlying the formation of these pores is technically challenging due to their requirement for specific receptors. Here, we tested a simple electrophysiological method enabling the real-time measurement of pore formation on model cells (Xenopus oocytes), which express the receptors of interest. We were thus able to elucidate the kinetics of pore formation, the limiting role of receptors in this process, and propose a complementary model to the standard model. We also demonstrated the ability of this method to detect pore formation of non-cognate pairs of subunits and suggest further applications to characterize pore-forming properties of other toxins, to identify new target receptors, or to screen therapeutic agents inhibiting the formation of pores.

## Introduction

*Staphylococcus aureus* represents an increasingly serious global health threat, due to the raising prevalence of methicillin-resistant *S. aureus* (MRSA) [1]. *S. aureus* is a ubiquitous Gram-positive bacterium, which is carried by the healthy population up to 20% persistently and 60% intermittently [2].

This pathogen can also cause a wide range of different infections, ranging from non-serious skin infections, to invasive, life-threating sepsis. The prevalence of MRSA infections in hospitals has become a major threat, due to the difficulty in treating multidrug-resistant strains in an already vulnerable population.

*S. aureus* releases a range of different virulence factors [3], such as pore-forming toxins, with expression varying depending on the site of infection [4] and growth conditions [5]. Among these factors are the bi-component leukocidins, which target different leukocytes, erythrocytes and endothelial cells [6] with high specificity relying on the type of receptors present in the membrane, and on which specific leukocidins bind to subsequently and drastically change their conformation and form cytotoxic transmembrane pores. In the current standard model, the S subunits are the components that recognize the host-receptors, and the F component [7] are required for the formation of the hetero-octameric pores [8]. In the nomenclature, the S subunits are cited first followed by the F subunits, with the known following pairs: the leukotoxins LukED and LukAB (also called LukGH), the gamma-hemolysins (Hlg) HlgAB and HlgBC and the Panton-Valentine

Leukocidins (PVL) LukS-PV and LukF-PV [9, 10]. Except for LukAB, the targeted receptors belong mainly to the GPCR family and the sub-families of chemokine and complement receptors, which have crucial roles in the innate and adaptive immune responses [11, 12]. The known target receptors are listed in reviews [13–15], including receptors used in this study that are CCR2 and ACKR1 targeted by HlgAB, C5aR1 targeted by HlgCB, and CCR5 targeted by LukED. Non cognate pairs are also reported as pore forming complexes such as LukE-HlgB targeting CCR5.

The architecture of the hetero-octameric pores comprises 4 S and 4 F subunits in alternate positions [16], but despite extensive research and the resolution of structures of the complex in different states (soluble, pre-pore and pore) [17], the exact molecular mechanisms of the formation of the complexes are not fully understood [13, 14, 16].

Most studies on the virulence of the various leukocidins rely on testing the permeability of cell membranes with nucleic acid dyes after exposure to the leukocidins, typically for 30 minutes or longer [11]. This is usually achieved by indirectly measuring the membrane permeability to organic dyes such as propidium iodide or others molecules that cannot easily transit through intact plasma membranes.

While the viability dyes are relevant for quantifying the cytotoxicity of leukocidins on various immune cells, they cannot discriminate between membrane permeability generated by pores and membrane permeability induced by cell death. To specifically study the mechanisms of pore formation at the molecular level, real-time measurements would be more appropriate. Electrophysiological techniques are powerful methods for structure-function studies of “pore-forming” proteins, mainly ion channels, allowing real-time recordings with a wide range of time scale (from milliseconds to minutes or hours). These techniques have been successfully used to record the pore formation of PFTs, such as the α-hemolysin, in artificial lipid bilayers [18]. Applying this approach to receptor-dependent PFTs, such as Hlgs, is less trivial since each targeted-receptor must be purified and re-constituted into artificial lipid bilayers that must preserve the native conformation and function of the receptors. Moreover, this process must be repeated for each measurement after pore incorporation in the lipid bilayer, which substantially limits the number of recordings.

Here, we present another approach that simplifies the functional studies of pore formation by the receptor-dependent leukocidins. This method has the advantages of not requiring receptor purification and insertion, to be operational in living cells, to allow rapid screening of various PFTs and receptors with electrophysiological robots, and to record in real-time the formation of pores in stages preceding cell lysis. The method is based on the heterologous expression of different PFT-targeted receptors in *Xenopus* oocytes [19]. Specific receptors are thus functionally expressed in individualized oocytes and currents generated by pores are recorded in real time with two-electrode voltage-clamp (TEVC) robots, HiClamp (Multichannel Systems) working in a 96-well plate format and requiring volumes of PFTs as low as ∼200 µL. Using this method, we explored the concentration effects of Hlg on the kinetics of pore formation, the impact of receptor expression levels and the sequential role of each subunit in the process of pore formation.

## Results

### Real-time recordings of HlgAB pore formation on CCR2-expressing Xenopus oocytes

To assess the capacity of the TEVC technique of detecting pore formation by receptor-dependent PFTs, the human CCR2 and CXCR4 chemokine receptors were heterologously expressed in *Xenopus laevis* oocytes (Fig 1) using mRNA micro-injection. After at least three days of incubation at 19 °C, real-time TEVC recordings were performed on individual oocytes during incubation in wells containing soluble PFTs. Initial attempts were made with two cognate soluble Hlgs (HlgA and HlgB) that target CCR2, while CXCR4 was used as a control.

**Fig 1.**
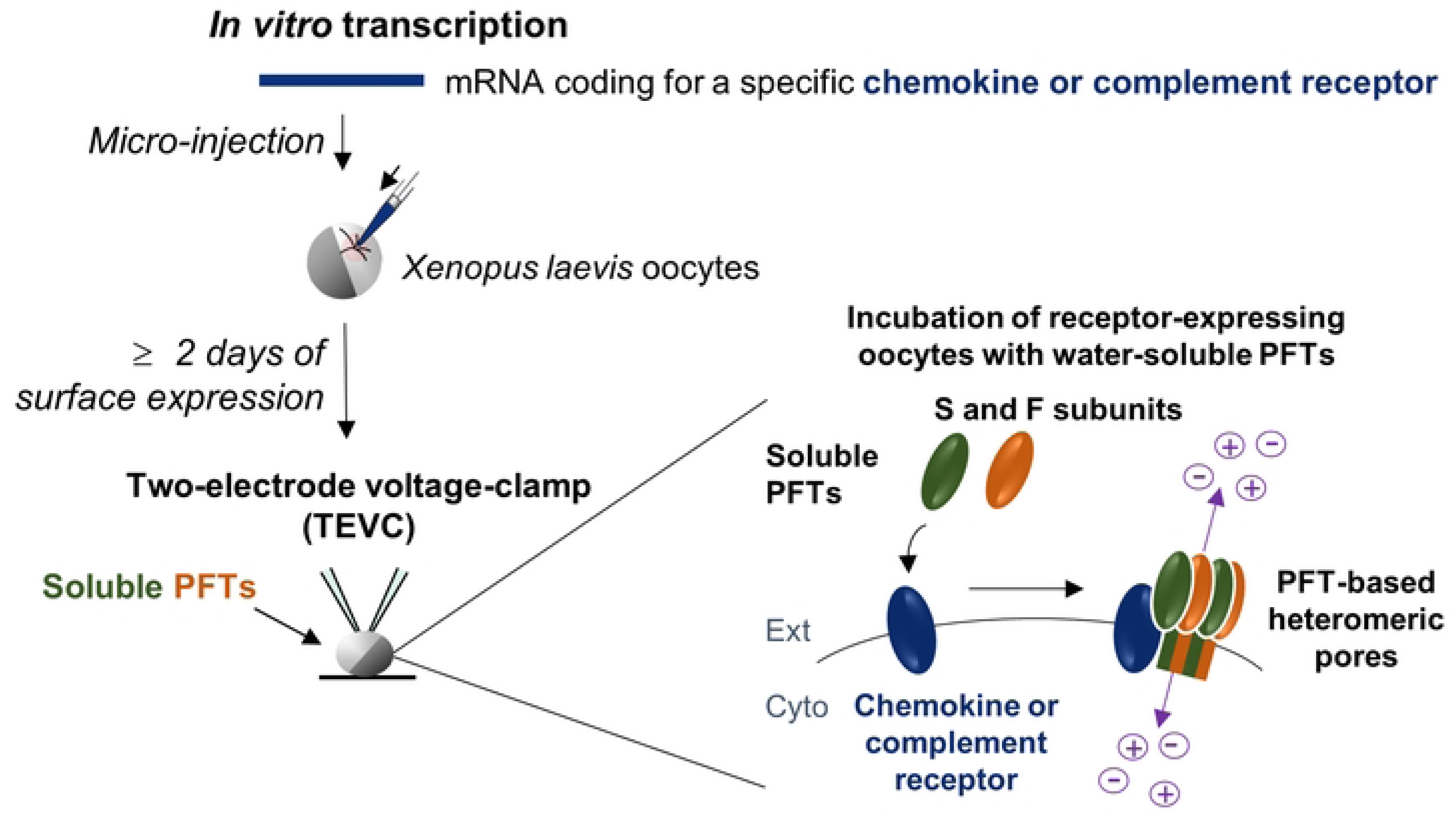
Method for expression and functional characterization of chemokine receptors heterologously expressed in *Xenopus laevis* oocytes. mRNAs are synthesized by *in vitro* transcription from receptor genes subcloned in a plasmid designed for overexpression in *Xenopus* oocytes. mRNAs are micro-injected into *Xenopus* oocytes in a volume of 50nl. After ≥ 2 days, receptors are significantly expressed in the plasma membrane of the oocytes. Using the two-electrode voltage-clamp technique, whole-cell currents are recorded in real time during applications of water-soluble pore forming toxins on the extracellular side of the cell.

HlgA and HlgB were first co-applied (noted HlgAB) on CCR2 or CXCR4 at a concentration of 1 µM of each subunit. Application of HlgAB generated large current amplitudes (−35.9 ± 3.7 µA, mean current at 1200 s) on CCR2-expressing oocytes (Fig 2A, red line), which is in agreement with reported CCR2-dependent pore formation of HlgAB [11]. The real-time recordings revealed an average delay of 38 s before the current increase occurred. This delay is consistent with a slow and progressive process. This process ended with a plateau in ∼500 s indicating a saturable mechanism. In control CXCR4-expressing oocytes, application of 1 µM of each subunit of HlgAB (Fig 2A, grey line) showed much lower current amplitudes (−5.1 ± 3.4 µA, mean current at 1200 s), which is also in agreement with the reported low potency of HlgAB on CXCR4 [11].

**Fig 2.**
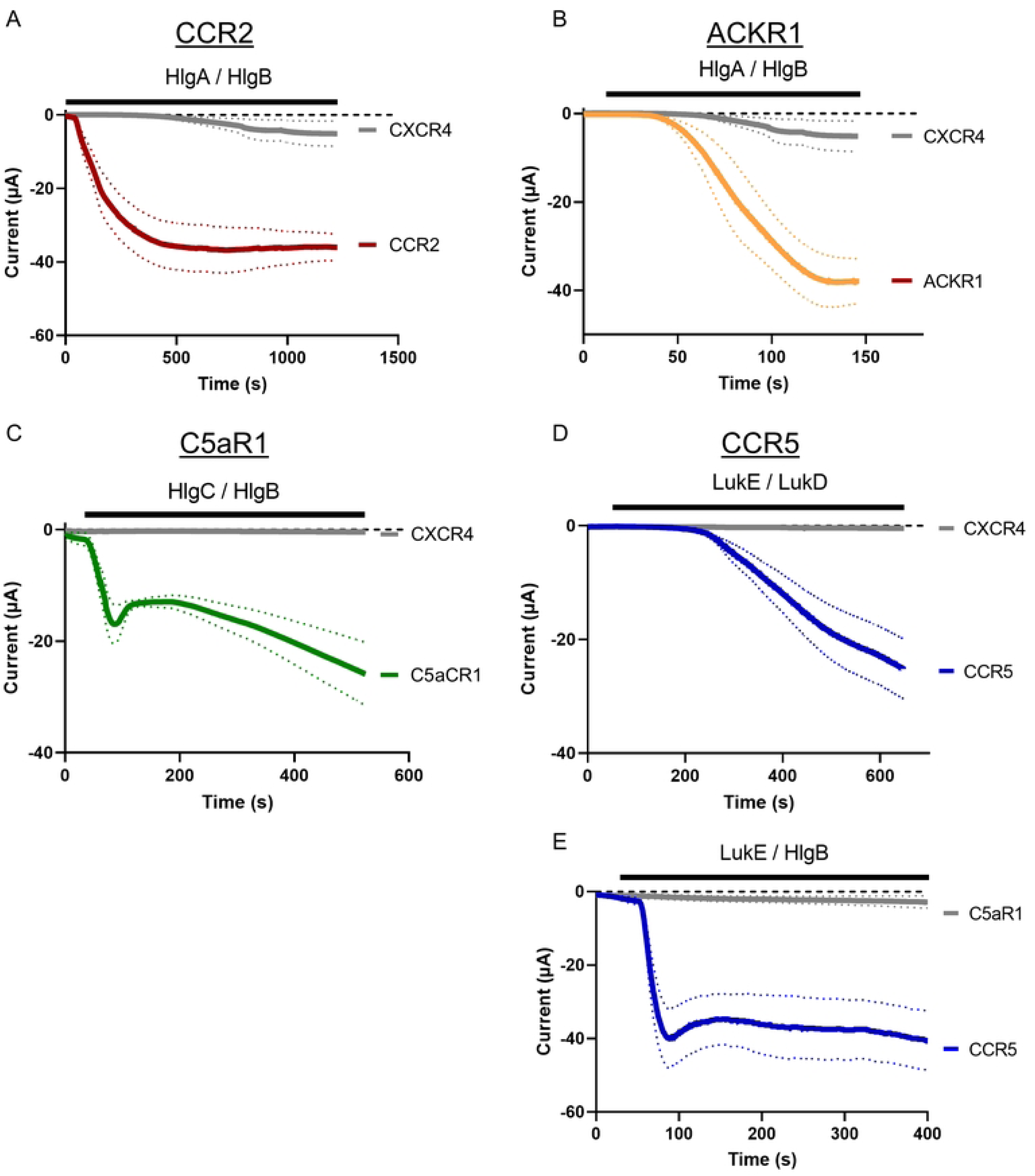
TEVC recordings of leucocidin-induced pore formation on *Xenopus* oocytes expressing different chemokine or complement receptors. **(A)** Whole-cell currents recorded by TEVC on oocytes expressing either CCR2 (red curve) and CXCR4 (gray curve, control) during incubation with HlgA/HlgB at 1 µM for each monomer. Curves are mean ± sem. n=18 for CCR2 and n=4 for CXCR4. The recording buffer is ND96 and the holding potential −50 mV. **(B)** Similar recordings on ACKR1-expressing oocytes (orange curve). n=6 for ACKR1 and n=4 for CXCR4. **(C)** Similar recordings on C5aR1-expressing oocytes (green curve) during incubation with HlgC/HlgB at 1 µM for each monomer. n=12 for C5aR1 and n=15 for CXCR4. **(D)** Similar recordings on CCR5-expressing oocytes (blue curve) during incubation with LukE/LukD at 1 µM for each monomer (n=5 for CCR5 and n=7 for CXCR4), or **(E)** during incubation with LukE/HlgB at 1 µM for each monomer as example of non-cognate pairing of leukocidins. n=5 for CCR5 and n= 4 for C5aR1.

### Extrapolation of the method to different chemokine receptors and leukocidins

To assess the versatility of the method, three other receptors were expressed in *Xenopus* oocytes: the human ACKR1, C5aR1 and CCR5. ACKR1, also called Duffy Antigen Receptor for Chemokines (DARC), is an atypical chemokine receptor that binds a large diversity of chemokines but does not activate G proteins [20]. C5aR1 is a complement receptor that binds the complement factor C5a [21] and CCR5 a chemokine receptor that binds various CCL chemokines [22], some of which are common to CCR2 that is phylogenetically very similar to CCR5 [23]. All three receptors (ACKR1, C5aR1 and CCR5) were challenged with their specific leukocidins: HlgAB, HlgCB and LukED, respectively, and the negative controls were performed on CXCR4. The TEVC results (Figs 2B-D) show the detection of pore formation on the three tested receptors in presence of their respective leukocidins, while no current or low currents with HlgAB were recorded in the negative controls with CXCR4, as expected. These results confirm the specificity of the responses and the ability of the technique to functionally characterize the formation of pores on different receptors and with different leukocidins.

On C5aR1, a “peak” of negative current amplitude is observed at ∼85 s and corresponds to the currents transiently generated by endogenous chloride channels (Cl_Ca_ or TMEM16A) activated by intracellular concentration of calcium [24]. These peaks are commonly observed when pores induce massive and rapid entry of calcium.

### Pore formation of non-cognate pairs of LukE/HlgB leukocidins on CCR5-expressing Xenopus oocytes

Apart from LukAB, the bi-component pore-forming leukocidins share over 60% sequence homology among S and F subunits. Most leukocidins can form functional pores using both cognate (LukED and HlgAB) and non-cognate combinations of subunits (e.g. LukE/HlgB and HlgA/LukD) [25, 26]. To test the ability of TEVC to detect the pore formation of non-cognate pairing of leukocidins, LukE/HlgB was applied on the known target CCR5 [13], and on C5aR1 as negative control (Fig 2E).

On CCR5, application of 1 µM of LukE and HlgB induced rapid (< 100 s) and large currents (−40.0 ± 8.0 µA at 88 s), while on C5aR1, the non-cognate pairs generated no significant currents (−2.8 ± 1.6 µA at 400 s versus −40.7 ± 8.0 µA on CCR5).

These results demonstrate that the TEVC is a useful and sensitive method to identify and characterize functional non-cognate pairs of PFTs on GPCRs.

### The concentration of HlgAB affects the kinetics of pore formation

To evaluate the sensitivity of the TEVC method for the detection of pore formation, HlgAB were co-applied at different concentrations from 0.1 nM to 1 µM of each subunit on CCR2-expressing oocytes (Fig 3A). Unexpectedly, only 1 µM of each subunit induced a significant increase of current (−42.2 ± 7.0 µA at 200 s), while previous binding studies by surface plasmon resonance indicated a high affinity of the human CCR2 for HlgA (K_D_ = 3.51 ± 0.29 nM), and membrane permeability assay with propidium iodide measured an EC_50_ of ∼2 nM of HlgAB on CCR2-expressing HEK cells in 30 min [11]. These previous results suggest that submicromolar concentrations of HlgAB should also form pores, but potentially with longer incubation times than the tested 200 s. Recordings in Fig 3B confirmed this assumption by measuring current increase with concentrations of HlgAB as low as 0.1 nM (each subunit) when applied over 1000 s and up to 2000 s. The traces revealed a kinetic of the pore formation that correlated with the concentration of toxins: the time of pore formation increases when the concentration decreases. The dashed line at 200 s on the Fig 3B, explains why only 1 µM of HlgAB generated currents in 200s in the Fig 3A.

**Fig 3.**
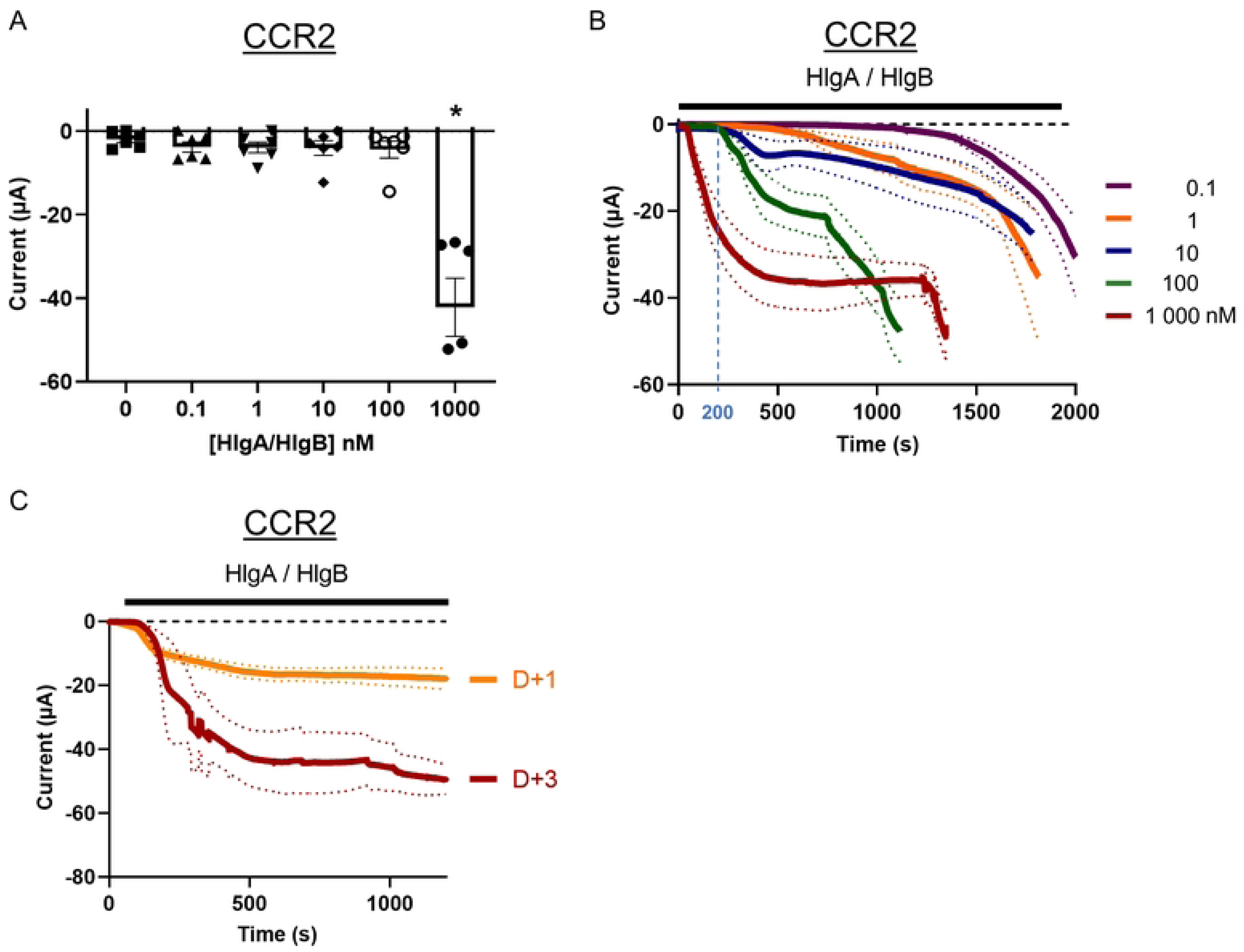
**Concentration-effect on the pore formation of HglA/HlgB on CCR2-expressing oocytes**. **(A**) Bar chart of currents at 200s induced by different concentrations (by monomer) of HlgA/HlgB on CCR2-expressing oocytes. TEVC recordings in ND96 buffer at a holding membrane potential of −50mV. n= 6, * P values=0,0158 for C=1000 nM (one-way ANOVA, ref=0 nM). **(B)** Time-course with extended time up to 2000s of currents induced by HlgA/HlgB at different concentration of each monomer: 0.1 nM (n=7), 1 nM (n=6), 10 nM (n=9), 100 nM (n=17), 1 µM (n=18). **(C)** Time-course of currents induced by HlgA/HlgB at 1 µM each monomer on CCR2-expression one day (D+1) or three days (D+3) post-microinjection of mRNA. n= 7 (D+1) and 4 (D+3). Bars and curves are Means +/- SEM.

Curves with concentrations of 10 nM and higher display biphasic profiles. In a first phase, current amplitudes increased up to a plateau in ∼500 s with amplitudes of −7.1 ± 3.2, −18.3 ± 4.9 µA and −35.8 ± 6.3 µA for 10, 100 and 1000 nM, respectively. In a second phase, the currents re-increased. For the highest concentrations (100 and 1000 nM), steep increase of current in the post-plateau phase resembles to massive leakage of the plasma membrane induced by cell death.

Consequently, the concentration of HlgAB significantly affects the kinetics of pore formation, which is in compliance with the oligomerization process leading to pore formation.

### The expression time of the CCR2 receptor limits the quantity of pores

The concentration of HlgAB affects the kinetics of pore formation, but the impact of the membrane receptor density is still unknown. To assess it, we recorded the pore formation at different times of CCR2 expression: one day (D+1) and three days (D+3) post mRNA micro-injection. D+1 is a suboptimal incubation time for surface expression of chemokine receptors under our experimental conditions [19]. The results (Fig 3C) showed that longer expression time of CCR2 had no significant effect on the kinetics of pore formation, but increased the current amplitude 2.7-fold (−42.7 ± 8.8 µA at D+3 *vs* −15.9 ± 2.0 µA at D+1 at 500 s). Since the concentration of HlgAB (1µM) is unchanged between the two conditions, the increase of current amplitude correlates with the increase of expression-time of CCR2. This result shows that receptors appear as a limiting factor for the formation of pores.

### Pores are formed by sequential applications of S and F subunits independently of the order

Based on the most common current models, the S subunit binds onto the receptor to recruit the F subunit and initiates the oligomerization process. However, previous studies demonstrated that some F subunits are also able to bind to the membrane of target cells independently of the S subunit and they participate in the efficacy of the oligomerization process. Their binding could occur either by interactions with lipids such as phosphatidylcholine [27] or membrane receptors. Indeed, it has been shown that LukF-PV not only binds onto the CD45 receptor, but it also leads to membrane permeabilization during sequential application with LukS-PV interspersed by a washing step [28].

Taking advantage of the real-time recordings by TEVC of pore-induced currents, we performed a similar approach of sequential applications of HlgAB on CCR2-expressing *Xenopus* oocytes by applying HlgA or HlgB first, and then the second subunit after a washing step (Fig 4A). In this configuration, Hlgs cannot create pre-formed heteromeric complexes. First applications of HlgA at 1µM (Fig 4B) did not generate changes in current amplitude, while the subsequent applications of HlgB after washing clearly generated an increase of current amplitudes with a calcium-induced peak of current of −21.4 ± 5.6 µA at 818 s, followed by a plateau of −8.9 ± 2.2 µA at 1175 s. This result shows the formation of pores when the two subunits are applied separately, indicating that preformed heterodimers are not necessary for pore formation. However, while the pore formation is fast as shown by the transient activation of Cl_Ca_ channels, the current amplitude is 1.7-times lower than the amplitude recorded during co-application of both subunits (−21.4 ± 5.6 µA for sequential applications *vs* −35.9 ± 3.7 µA for co-applications). This result indicates that a factor limits the formation of pores during sequential applications of HlgA and HlgB. The recordings being performed at D+4 post-micro-injection, the density of CCR2 on the cell surface was optimal, and the concentration of both subunits was identical between co- and sequential applications (1 µM each). Consequently, the limiting factor would be the missing water-soluble subunit (HlgA) during the sequential application of HlgB.

**Fig 4:**
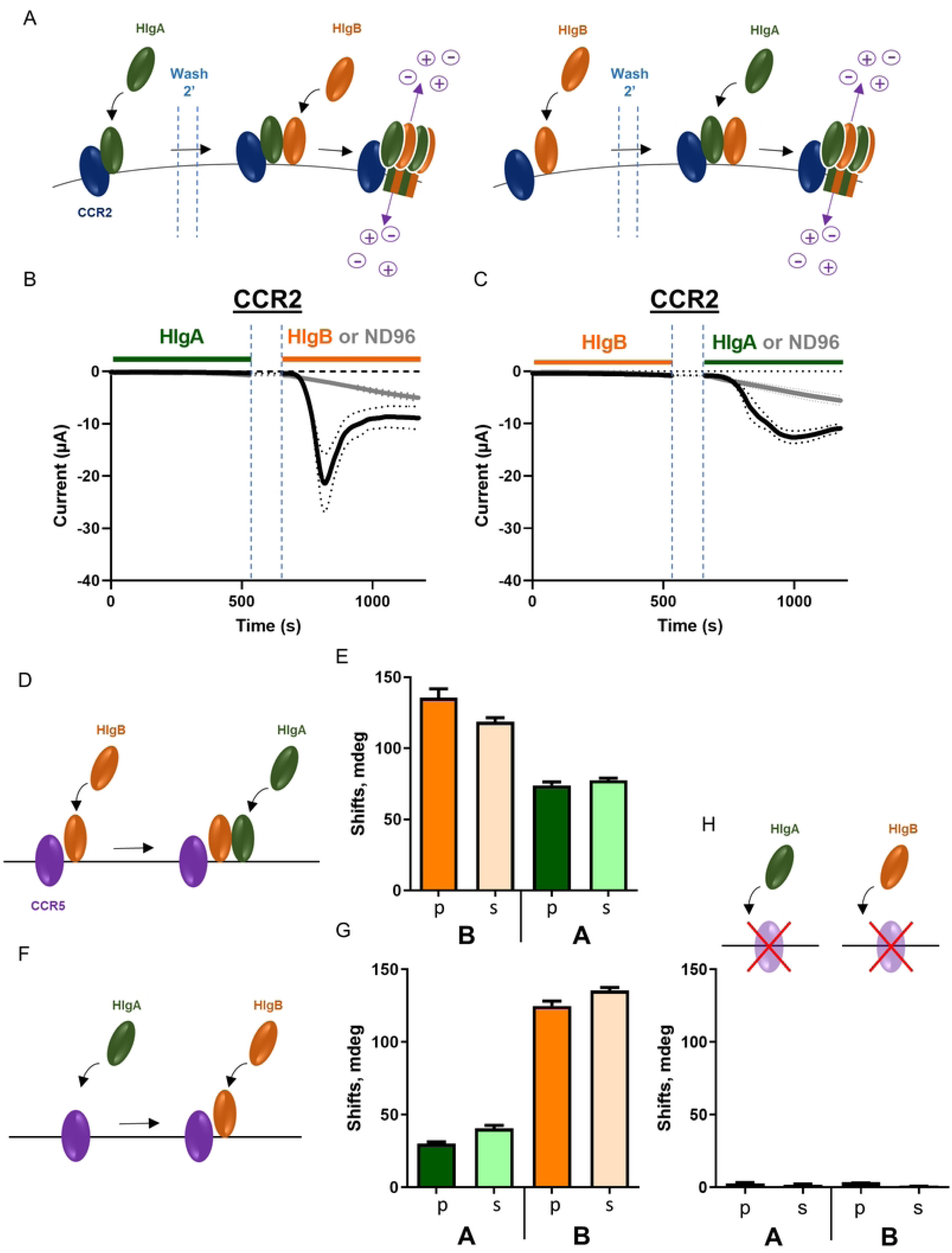
Sequential applications of soluble PFTs enable pore formation. **(A)** Schemes of sequential applications at 1 µM each subunit of either HlgA and then HlgB (left panel), or the reverse sequence (right panel) interspersed by 2 min washing in a flow of buffer ND96. **(B-C)** TEVC recordings of sequential applications depicted in panel A with either application of the second subunit after washing (black curves) or application of ND96 buffer as negative control (grey curves). n=6 (HlgA-HlgB), n=12 (HlgA-ND96), n=12 (HlgB-HlgA), n=10 (HlgB-ND96). Curves are Means +/- SEM. **(D)** Scheme of sequential application of HlgB and then HlgA on CCR5 immobilized in a lipid bilayer on the surface of PWR prism. **(E)** Means +/- SEM of change in resonance angles (Shifts) for (p- and s-polarisation) during HlgB and the HlgA applications as depicted in panel D. **(F-G)** similar panels as D and E respectively, with application of HlgA first and then HlgB. **(H)** Negative controls with application of HlgA or HlgB on surfaces with a lipid bilayer exempt of CCR5.

The opposite sequence was tested by applying HlgB first, then HlgA (Fig 4C), and pores were also formed during application of the second subunit with an amplitude at the end of the recording in the same range as in Fig 4B (−10.9 ± 0.8 µM at 1175 s). This result suggests that some HlgB subunits are present on the oocyte surface and form pores with HlgA during its sequential application. This is in agreement with previous observations of interaction of HlgB with the membrane of erythrocytes [29], and the formation of heteromeric pores with pre-bound LukF-PV [28].

### HlgB also binds to the CCR5 chemokine receptor

The formation of pores by sequential applications of HlgB and then HlgA suggests that HlgB subunits bind onto the surface of oocytes.

HlgB has the particularity to form cognate and non-cognate pairs with all leukocidins (except for LukAB) and form pores in the presence of diverse receptors (C5aR1, C5aR2, CCR5, CXCR1, CXCR2, CCR2 and ACKR1) [13]. Binding of HlgB on a receptor has been observed on ACKR1 by native mass spectrometry and confirmed by time-resolved fluorescence energy transfer (TR-FRET) and bioluminescent resonance energy transfer (BRET) [30]. To confirm the ability of HlgB to bind to another chemokine receptor, we took advantage of the previous immobilization of purified CCR5 receptors in planar lipid membranes on the surface plasmon waveguide resonance (PWR) sensor [31],[32]. PWR is a method based on surface plasmon resonance for investigations of molecular interactions that, due to the presence of a dielectric surface (silica) on top of the plasmon resonance layer (silver), allows the use of both *p*- and *s*-polarized light to create resonances. This important feature when applied to oriented thin films as proteolipid membranes provides in addition to mass changes, information on anisotropy [33],[34]. In sequential applications to a proteolipid membrane containing the CCR5 receptor deposited on the sensor surface, the addition of HlgB first (Figs 4D and E) shows a clear increase in both resonance angles polarisation (*p-* and *s-*pol) that reflects a large mass gain as a consequence of membrane binding. The change is slightly anisotropic with *p*-shifts being higher than *s*-shifts (p) and change of anisotropy (s). On the other hand, the application of HlgA first (Figs 4F and G) shows much lower changes of both *p*- and *s*-shifts (lower mass gain), which was expected since CCR5 is not reported as a target of this S subunit, but suggests weak interactions with the receptor. Application of HlgB or HlgA on membrane without CCR5 does not induce *p-* and *s-*shifts (Fig 4H), confirming the absence of non-specific binding on the lipids. These results show that HlgB interacts with CCR5 and the binding is anisotropic in nature due either to the intrinsic anisotropy of the protein itself or to changes in the receptor-containing bilayer, or both.

## Discussion

### TEVC is a suitable method for the functional characterization of receptor-dependent PFTs

At the molecular scale, the functional characterization of pore formation by GPCR-dependent PFTs is technically challenging due to the need to express and isolate specific chemokine and complement receptors in reconstituted systems. The method described in the present article facilitates the functional characterization of PFT-formed pores by enabling heterologous expression of a wide variety of membrane receptors without delicate, time-consuming purification and reconstitution in a lipid bilayer of each receptor. The present study demonstrates that the TEVC technique is a suitable method for monitoring pore formation on individual living cells (*Xenopus* oocytes) expressing specific GPCRs. This approach provides qualitative and quantitative information in early phases of pore formation with high sensitivity and in stages preceding the cytotoxic effect of the toxins. Another advantage of this approach is the ability to rapidly screen and characterize both PFTs and membrane receptors using both automated mRNA micro-injection (RoboInject, MCS) and TEVC recordings (HiClamp, MCS).

This method could be used in structure-function studies of PFT variants having divergent receptor tropisms in different bacterial lineages [35, 36], or in studies cataloguing the diversity of toxins having pore forming activities [10], even in other genus than *Staphylococcus* [37]: e.g. HlyA from *E. coli* targeting the 6-cyano-7-nitroquinoxaline-2,3-dione receptor [38] and pneumolysin from *S. pneumoniae* targeting the mannose receptor C type 1 [39].

The technique is theoretically not restricted to GPCRs and could also be extended the repertoire of PFT targets from other membrane protein families. Thus, different examples of non GPCR PFT targets have also been reported [40] such as disintegrin and metalloproteinases ADAM10 for the α-hemolysin (also able to form pores without receptors) [41], CD59 intermedilysin for the streptolysin O [42], CD45 receptor protein tyrosine phosphatase for the LukF-PV subunit [28] and the heteromeric complex CD11b/CD18 (αM/β2) integrin [43] and HVCN1 ion channel [6] for LukAB.

The electrophysiological robots are also suitable to screening approaches and could be used to identify PFT inhibitors based on various pharmacological strategies [4, 44] including engineered non-pore forming toxins [45] or receptor ligands [46]. In anti-cancer therapies, engineered PFTs targeting specific receptors present on the surface of cancerous cells could be also functionally characterized by TEVC for developing synthetic biology approaches [47].

The limitations of this approach are: 1) the requirement of sufficient surface expression of receptors in *Xenopus* oocytes; 2) the purification of water-soluble PFTs; 3) the use of non-endogenous cells that are also not optimal for biochemistry, optical and fluorescence microscopy and cytotoxic assays; and 4) the cost of the robots. Moreover, the TEVC method records macroscopic currents and consequently lacks the single pore resolution required for determining unitary conductance, open probability or pore current heterogeneity. *Xenopus* oocytes also express endogenous Cl_Ca_ channels that partially and transiently increase the recorded kinetics of pore-formation.

The method has also advantages such as the possibility: (1) to express a high number of different receptors without purification and membrane insertion; (2) to screen receptors and PFTs using HiClamp robots; (3) to maintain natural and alive cellular environment with a natural asymmetry of the lipid bilayer; (4) to measure in real time the formation of pores; (5) to increase sensitivity thanks to the large amount of ions Na^+^, K^+^, Cl^-^, Ca^2+^ crossing the pore magnified by the endogenous Cl_Ca_ channels in *Xenopus* oocytes; and (6) to perform experiments with inexpensive and non-hazardous solutions.

### Kinetics of pore formation

The TEVC technique enables PFT studies recordings in real time, allowing characterization of toxin concentrations on the kinetics of pore formation. Increasing concentrations of HlgAB induced a reduction of the delay (T_Lag_) preceding the pore formation from ∼1100 s at a subnanomolar concentration (0.1 nM each subunit) to ∼40 s at 1 µM. This concentration effect on T_Lag_ was also observed on real-time measurements of erythrocytes swelling in presence of aerolysins [48]. While aerolysins are receptor-independent PFTs, the pore formation process follows similar steps such as diffusion of soluble monomers, binding on the cell surface, conformational changes, oligomerization and pore formation that are related to the concentration of monomers [49]. However, some differences are reported depending on the PFTs. For example, heptameric α-hemolysin shows rapid (<5ms) apparent-single step pore formation [50], suilysin forms transient hemi-pores by in-membrane oligomerization [51], while Hlgs (HlgAB) bind to receptors, dimerize with conformational changes (release of amino-latch and stem domains), oligomerize with alternate positions of subunits and form pre-pore followed by the transmembrane pore insertion [17, 52].

In this later process, some questions remain open, notably on the role of the receptors in the oligomerization process. Our results with different incubation time of microinjected oocytes (Fig 3C) suggest that the quantity of receptors available at the cell surface is a limiting factor for pore formation in a saturated concentration of toxins. Since no additional pores are formed in stationary phase (during our recording period of 1200 s), it suggests that free soluble subunits are unable to form pores with saturated receptors once the pores are formed. This receptor saturation effect is supported by observations on polymorphonuclear (PMN) cells that showed an increase of LukAB cytotoxicity when the membrane receptor density increased in activated-state [53]. The main hypothesis for lack of functional neo-pore formation is a stable interaction of formed-pores with receptors. However, partial dissociation of leukocidin pores (LukSF-PV) has been described on the complement receptor C5aR [54]. Consequently, potential dissociation of HlgAB pores from the chemokine receptor CCR2 cannot be excluded under physiological conditions and in other timescales.

### Model of sequential formation of pores

The pore formation by leukotoxins is a dynamic process involving receptor binding and oligomerization with alternate positions of the two PFT subunits until the formation of an octameric complex [10, 13, 15, 17, 40]. Two scenarios could explain this organization. The first one is inspired by LukAB [55], which has the particularity of forming constitutive water-soluble dimers. Consequently, LukAB forms alternate subunit complexes by recruitment of heterodimers [6]. A similar scenario could occur for Hlgs since water-soluble HlgAB heterodimers have been observed by native mass spectrometry [30]. However, their relative abundance is low compared to monomers. A second scenario is preferred in which, F subunits are recruited by S subunits bound to receptors. Binding of the S subunit onto the receptor would unlatch its stem domain, promoting the oligomerization of S and F subunits that unlatch their stem domains until the final formation of the functional β-barrel transmembrane pore [6].

The TEVC recordings with sequential applications of HlgA and then HlgB, interspersed by a washing step (Fig 4B), showed pore formation during incubation in HlgB which indicates the ability of monomers to oligomerize until the formation of hetero-octameric pores. However, the current amplitude in sequential application is lower than in simultaneous application of both subunits. The limiting factor during these sequential applications is the absence of water-soluble HlgA during HlgB application. Consequently, the lower current amplitude observed during sequential applications suggest a role of the unbound S subunit in the formation of pores. It is not possible to discriminate between a lower amplitude due to the formation of partial pores lacking protomers of HlgA or the formation of a limited number of complete octameric pores made by receptor-bound HlgA. However, functional and structural evidences, such as TEVC recordings showing surface expression of CCR2 as a limiting factor for pore formation (Fig 3C) and the formation of octameric pre-pore before transmembrane pore formation [17], are in favor of the second hypothesis of a limited number of complete octameric pores during sequential applications of HlgA and HlgB. Moreover, crystal structures of LukAB bound to the receptor binding site moiety (CD11b-I), show an interaction of CD11b-I with adjacent protomers, suggesting that CD11b is involved in both binding of the leukotoxins and the oligomerization process. Thus, the receptors would play a role during the formation of heteromeric pores with alternate positions of S and F subunits [55].

### The F subunit also binds to the cell surface to form heteromeric pores

Sequential application of Hlgs has also been performed in reverse order with the F subunit (HlgB) first followed by the S subunit (HlgA) with interspersed washing step (Fig 4C). According to the model of recruitment of F subunits by S subunits bound to receptors, no pores should be observed. However, TEVC recordings clearly showed pore formation during the incubation in HlgA, indicating that subunits of HlgB remained bound on the cell surface and oligomerize with newly added HlgAs. This result is supported by previous reports of F subunit binding to the cell surface such as LukD and HlgB bound onto the membrane of erythrocytes [29]. Moreover, LukF-PV not only binds onto CD45, but also leads to membrane permeabilization of murine C5aR1^KI^ neutrophils during sequential application with the S subunit (LukS-PV) interspersed with a washing step [28]. On that basis, an alternative model of pore formation by bi-component leukotoxins is proposed with simultaneous interactions of S and F subunits of PVL leukotoxins with their receptors C5aR1 and CD45, respectively [56].

However, CD45 is not a receptor of HlgB [28], which raises the question of its site of interaction with the membrane [4]. It could be either lipids such as phosphatidylcholine [57, 58], “co-receptors” like CD45 for LukF-PV[28], a mix of lipids, other membrane components or the chemokine receptor.

Results from PWR with immobilized CCR5 (Figs 4D-G), showed binding of HlgB on membranes containing CCR5, which is a target of the LukE-HlgB non-cognate pairing leukotoxin (Fig 2E). Native mass spectrometry and BRET also showed binding of HlgB on the purified chemokine receptor ACKR1 with a 1:1 ratio [30]. These *in vitro* experiments suggest that HlgB could interact with different chemokine receptors.

### Hybrid model of pore formation by sequential process and receptor clustering

The molecular mechanism of oligomerization process of S and F subunits is not precisely known. The standard model is based on sequential interaction of the two subunits starting from interaction of a S subunit with its receptor (Fig 5A). However, several lines of evidence indicate that the F subunits can also interact with the targeted cell surface either through lipids or membrane receptors. Sequential applications of isolated subunits indicate that S and F subunits can adsorb on the cell surface and form pores when the second subunit appears. This mechanism could appear when the concentration of the subunits is low, such as in the early stages of infection, and would improve the efficacy of monomeric subunits to form heteromeric pores by accumulating S and F subunits onto the cell surface and by progressively forming pores when the second subunits reach the cell surface. This mechanism could progressively evolve into a hybrid mechanism involving oligomerization of both water-soluble and membrane bound subunits (Fig 5B), increasing the kinetics of pore formation. Finally, when the concentration of both subunits is optimal, the standard model which is the most efficient model

**Fig 5:**
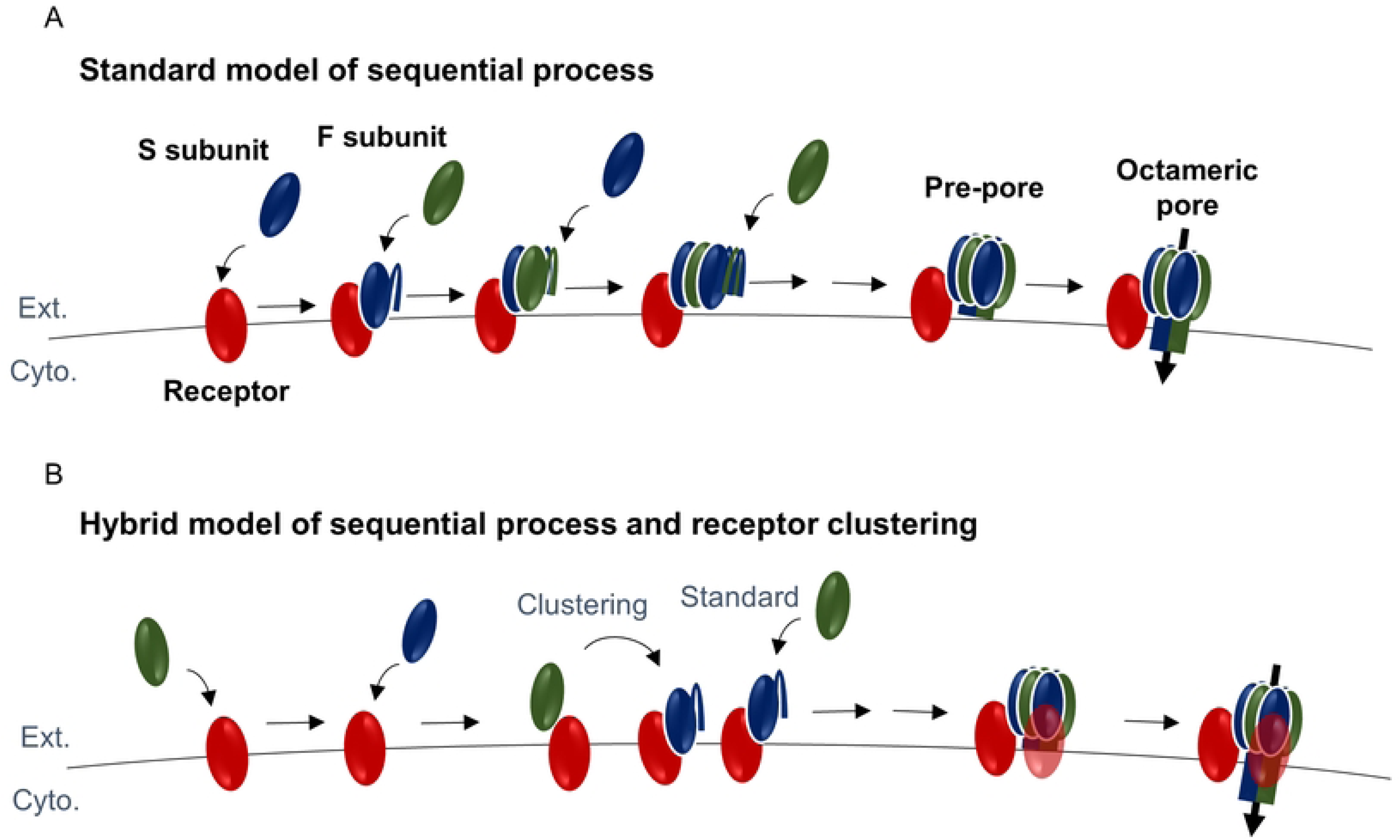
Standard and hybrid models of pore formation by leukocidins . **(A)** standard model of leucocidin pore formation by sequential process. Binding of the S subunit onto its target receptor unlatches the stem domain and triggers alternate oligomerization of S and F subunits until the formation of an octameric prepore followed by the formation of the pore by insertion of the stem domain β-barrel into the lipid bilayer. **(B)** Hybrid model based on the standard model supplemented with a process of clustering of the S and F subunits bound on receptors at the membrane surface. Both models could co-exist and evolve with the quantity of toxins produced at different stages of infection.

Overall, the oligomerization mechanism would not be based on a single scenario but on complementary mechanisms depending on the concentration of circulating subunits.

## Materials and Methods

### Molecular biology

All human chemokine and complement receptors (CCR2, CXCR4, CCR5, C5aR1 and ACKR1) genes were subcloned in pGH or pXOOM [59] vectors, designed with 5’ and 3’ untranslated regions (UTRs) of *Xenopus* β-globin for overexpression of proteins. CCR2 is from the B isoform (uniprot# P41597-2). mRNA was obtained by *in vitro* transcription using the T7 mMessage mMachine Kit (ThermoFisher Scientific) and purified using the standard phenol:chloroform protocol. Quality control was done by electrophoresis profile and the RNA was quantified using nanodrop spectrophotometer.

### Expression and purification of leukocidins

M15 *E. coli* strains (Qiagen, Germany) were transformed with the expression plasmids including HlgA or HlgB. These transformed strains were cultured in 2 L of Luria-Bertani broth (LB) supplemented with 100 μg/mL ampicillin and 30 μg/mL kanamycin at 37°C under 150 rpm agitation until reaching an optical density (OD) of 0.3. Prior to induction, the culture was pre-cooled at 15°C for 30 minutes under 150 RPM agitation at 15°C. Isopropyl β-D-1-thiogalactopyranoside (IPTG) was then added to a final concentration of 0.5 mM and the cultures were grown overnight. The induced cultures were centrifuged at 4500 g for 15 minutes and the resulting pellet was resuspended in 40 ml of lysis buffer (20 mM HEPES (pH 7.5), 150 mM NaCl, 50 mM imidazole, 1 tablet of cOmplete EDTA-free protease inhibitor (Roche), 10 μg/ml DNase I, and 10 μg/ml RNase). The cells were lysed by subjecting them to four cycles using the microfluidizer LM20 (Microfluidics, USA) at 17000 psi while maintaining a temperature of 4°C. Subsequently, the lysate was clarified by centrifugation at 38500 g or relative centrifugal force (RCF) for 20 minutes at 4°C. His-tagged proteins were purified using immobilized metal chelate affinity chromatography on a 1 ml His-Trap HP column (Cytiva). Prior to loading the lysate, the column was equilibrated with 20 mM HEPES (pH 7.5), 150 mM NaCl, and 50 mM imidazole. A stepwise elution was performed using elution buffers (20 mM HEPES (pH 7.5), 150 mM NaCl, and 250 mM or 300 mM imidazole). Following purification, the eluate from the final elution step was dialyzed against the buffer containing 20 mM HEPES (pH 7.5) and 150 mM NaCl overnight at 4°C using a Spectra/Por 4 dialysis membrane.

Other leukotoxins were expressed with (His)6-tags at their carboxyl termini in competent C43 (DE3) *Escherichia coli* cells (New England Biolabs) for HlgA and in BL21 (DE3) *E. coli* cells (New England Biolabs) for HlgB. Transformed cells were grown at 37 °C in Terrific broth for HlgA and in Luria–Bertani broth for HlgB supplemented with 100 mg/mL ampicillin to an optical density of 0.6. Expression was then induced at 22 °C by addition of 0.5 mM isopropyl ß-D-1-thiogalactopyranoside. Cells were harvested by centrifugation (3,000 rpm), and cell pellets were stored at 80 °C until purification. After thawing the frozen cell pellets, cells were lysed by sonication in a lysis buffer consisting of 20 mM Tris (pH 8), 300 mM NaCl, 2 mg/mL iodoacetamide (Sigma-Aldrich), and protease inhibitors (Leupeptin [Euromedex], Benzamidine, and PMSF [Sigma-Aldrich]). Lysed cells were centrifuged (16,000 rpm), and the supernatant was adjusted to 40 mM imidazole and loaded onto a nickel nitrilotriacetic acid agarose resin. The resin was washed with 10 CV wash buffer 1 consisting of 50 mM Hepes (pH 7.5) and 1 M NaCl and with 10 CV wash buffer 2 consisting of 50 mM Hepes (pH 7.5) and 150 mM NaCl supplemented with 40 mM imidazole. Bound (His)6-toxins were eluted with wash buffer 2 supplemented with 200 mM imidazole. The eluted solution of toxins was concentrated to 500 μL using 30-kDa spin filters (Millipore) and further purified by SEC on a Superdex 200 Increase 10/300 column (GE Healthcare) in 50 mM Hepes (pH 7.5) and 150 mM NaCl. Fractions containing monodisperse toxins were collected and concentrated

### Heterologous expression in *Xenopus* oocytes

Expression of chemokine receptors in *Xenopus* oocytes has already been described elsewhere [19]. Briefly, *Xenopus* oocytes were defolliculated using a treatment with type Ia collagenase at 2mg/ml in Buffer A (88mM NaCl, 1mM KCl, 0.82mM MgSO_4_.7H_2_O, 2.4mM NaHCO_3_, 16mM HEPES, pH 7.4). Healthy oocytes in stages V and VI were manually selected and stored in modified Barth’s solution (Buffer A + 0.41 mM CaCl_2_ + 0.3 mM Ca(NO_3_)_2_.4H_2_O). Then 50 nl of mRNA containing 2 ng of receptor-coding RNA in DEPC-treated milliQ water, were injected in oocytes using either a microinjector Nanoject (Drummond) device or the RoboInject (Multichannel Systems). Oocytes were incubated in 96-wells plate with modified Barth’s solution + antibiotics (penicillin 100U/ml, streptomycin 0.1mg/ml, gentamycin 0.1mg/ml) for at least 48h (unless specified) at 19°C. Because of the variability in expression, experiments were done on several oocytes from the same and different batches of oocytes. Animals were housed and bred at CEA (Agreement # D 38 185 10 001) and currently in the « Plateforme de Haute Technologie Animale (PHTA) » UGA core facility hTAG, Inserm US46, CNRS URA2019 (La Tronche, France), EU0197, Agreement D38-516 10 006. Animal housing and procedures were conducted in accordance with the recommendations from the Direction des Services Vétérinaires, Ministry of Agriculture of France, according to European Communities Council Directive 2010/63/EU and according to recommendations for health monitoring from the Federation of European Laboratory Animal Science Associations. Protocols involving animals were reviewed by the CEA ethic committee and approved by the Ministry of Research (APAFIS#30915-2021040615209331 v1 to CJM).

### Electrophysiological recordings

Whole-cell currents were recorded with the automated two-electrode voltage clamp (TEVC) technique (Robot HiClamp, Multichannel Systems). Microelectrodes were filled with 3M KCl and oocytes were bath in ND96 solution (91 mM NaCl, 2 mM KCl, 1.8 mM CaCl_2_, 1 mM MgCl_2_, 5mM HEPES, pH 7.4). The TEVC voltage value was held at −50 mV. Average values are presented as mean ± SEM. Soluble PFTs were applied with single concentrations in wells of the ’ligand plate’ of the TEVC robot, or with increasing concentration for concentration-effect recordings. For sequential applications of subunits, the following recording protocol has been applied: 15s in a well with ND96, 500s in a well with the 1^st^ Hlg, 120s of washing in a stream of ND96, 15s in a well with ND96 and 500s in a well with the 2^nd^ Hlg.

### Plasmon waveguide resonance measurements

Plasmon waveguide resonance (PWR) measurements were performed in a homemade instrument equipped with a He-Ne laser, with a fixed wavelength in the visible region (λ=632 nm; Melles Griot), a rotating table allowing the incident angle to be changed by steps of ≤1 mdeg (thus resolution being on that order; Newport) and a photodiode detector (Hamamatsu) to measure the reflected light as a function of the incident angle. The polarization angle of the incident beam is placed at 45° allowing both *p*- and *s*-polarized spectra to be obtained within the same angular scan. The sensor consisted in a BK-7 prism coated with silver and silica (to support waveguide modes) [60]. Full spectra, minimum resonance position and spectral depth is acquired in real time via a Matlab interface. More detailed information on the principles of the method can be found in [33]. All measurements were performed at 22 °C in a temperature-controlled room under dim light. 50 μL solution of proteoliposomes composed of egg PC, cholesterol and CCR5 provided by Synthelis Biotech were applied to the PWR sensor and incubated for 15 min. Following that the cell sample in contact with the sensor was filled with TBS buffer and spectra recording (previous to that spectra of the buffer alone are recorded as reference). When the PWR signal was stable (no changes in the resonance position), HlgA or HlgB was added in incremental fashion up to saturation and global shifts in both *p*- and *s*-pol recorded. A control experiment was performed to investigate HlgA and HlgB binding to membranes lacking CCR5. Despite using the same amount of proteoliposomes, some heterogeneity in membrane deposition and level of CCR5 present can occur which can impact the interaction of HlgA and HlgB. To correct for that, the data was normalized by taking into account the spectral shifts difference observed upon proteoliposome deposition (relative to buffer alone).

## Acknowledgments

We thank Hervé Pointu, Soumalamaya Bama Toupet and Charlène Caloud from CEA for the management and the maintenance of *Xenopus* at We thank Candice Hackney, Emeline Mercier, Maryline Cossin, Hervé Lerat, Nadia Hassani, Bertrand Favier and other personnels of PHTA facility for animal housing and care. We thank Bruno Tillier and Synthelis Biotech for providing customized CCR5-proteoliposome to IDA. We thank Gérard Lina and Francois Vandenesch for intial work with AMDG during the ANR project PVLSuscept. IBS acknowledges integration into the Interdisciplinary Research Institute of Grenoble (IRIG, CEA).

## Author Contributions

LL: investigation, formal analysis, writing original draft; SH: investigation, formal analysis, writing original draft; GA: investigation, formal analysis; LB: resources; JDS: investigation, formal analysis; CMG: resources; SG: resources; JM: resources; NV: resources, manuscript revision; TV: resources; IDA investigation, formal analysis, writing original draft; AMDG: conceptualization; CJM: conceptualization, supervision, investigation, formal analysis, writing review & editing.

